# Microscope-AOtools: A generalised adaptive optics implementation

**DOI:** 10.1101/2020.06.18.158972

**Authors:** Nicholas Hall, Josh Titlow, Martin J. Booth, Ian M. Dobbie

**Affiliations:** Micron Advanced Bioimaging Unit, Department of Biochemistry, University of Oxford, South Parks Road, Oxford, OX1 3QU, United Kingdom; Davis Lab, Department of Biochemistry, University of Oxford, South Parks Road, Oxford, OX1 3QU, United Kingdom; Department of Engineering Science, University of Oxford, Parks Road, Oxford OX1 3PJ, UK

**Keywords:** Adaptive Optics, Software, Microscopy, Aberration Correction

## Abstract

Microscope-AOtools is a software package which allows for a simple, robust and generalised implementation of adaptive optics (AO) elements. It contains all the necessary methods for set-up, calibration, and aberration correction which are simple to use and function in a robust manner. Aberrations arising from sources such as sample heterogeneity and refractive index mismatches are constant problems in biological imaging. These aberrations reduce image quality and the achievable depth of imaging, particularly in super-resolution microscopy techniques. AO technology has been proven to be effective in correcting for these aberrations and thereby improving the image quality. However, it has not been widely adopted by the biological imaging community due, in part, to difficulty in set-up and operation of AO, particularly by non-specialist users. Microscope-AOtools offers a robust, easy-to-use implementation of the essential methods for set-up and use of AO techniques. These methods are constructed in a generalised manner that can utilise a range of adaptive optics elements, wavefront sensing techniques and sensorless AO correction methods. Furthermore, the methods are designed to be easily extensible as new techniques arise, leading to a streamlined pipeline for new AO technology and techniques to be adopted by the wider microscopy community.

## 0.1 Introduction

Many of the recent innovations in biological imaging have revolved around the quest for greater resolving power, ultimately culminating in the advent of super-resolution microscopy techniques. However, there is often a difference between the theoretical resolution and the practical resolution obtained in biological imaging. This is particularly true for live, thick samples, which are interesting to biological researchers for their ability to show dynamic biological processes *in situ*. How close the theoretical and practical resolutions are to one another is largely dependent on the optical aberrations present, most of which arise from the heterogeneity of the biological sample itself [1, 2]. These aberrations compromise image quality, decreasing contrast and resolution, by distorting the optical wavefront[3, 4]. Implementing adaptive optics (AO) in microscopy has already been shown to be highly effective at reducing these aberrations and yielding significant improvements to image quality[5, 6]. The widespread use of AO in microscopy would therefore be a significant boon to biological research.

Unfortunately, whilst multiple proof of principle systems in AO microscopy have been demonstrated, use of AO has yet to be widely adopted[7, 8, 9, 10]. This is due, in large part, to the complicated nature of measuring the wavefront deformations (and therefore the aberrations) present in a sample. While methods for directly measuring the wavefront do exist, they carry additional complications such as limiting what can be images for correction e.g. only point sources[11]. Therefore indirect wavefront sensing, or sensorless AO, is generally preferred[12]. In sensorless AO methods, some quality of the sample images, such as contrast or spatial frequency content, is evaluated and this quality is maximised by varying the wavefront deformation applied to the corrective element. Most proofs of principle AO microscopes implement a single method of correction that is particularly suited to the specific imaging modality and/or sample being used. So far, a robust, generalised, easy-to-use implementation which incorporates multiple AO methods for multiple sample types and imaging modalities has yet to be presented[13].

Microscope-AOtools provides such a generalised solution. It utilises Python-Microscope, an open-source hardware control software package, to provide control over the physical hardware necessary for AO implementations. It incorporates methods for calibrating an AO element, evaluating the success of the calibration in recreating aberrations and performing both direct wavefront sensing and, so called, sensorless adaptive optics corrections where aberrations are detected directly from images. The methods for sensorless AO correction can utilise a number of different image quality metrics. The methods presented are built in such a manner to enable easy switching between different AO elements, wavefront sensing techniques and image quality metrics. They are designed to be easily extensible so that new technology and techniques can be readily incorporated.

## 0.2 Principles behind Microscope-AOtools

Designing an AO enabled system follows a predicable workflow outlined in Figure 1 consisting of four phases:

**Figure 1:**
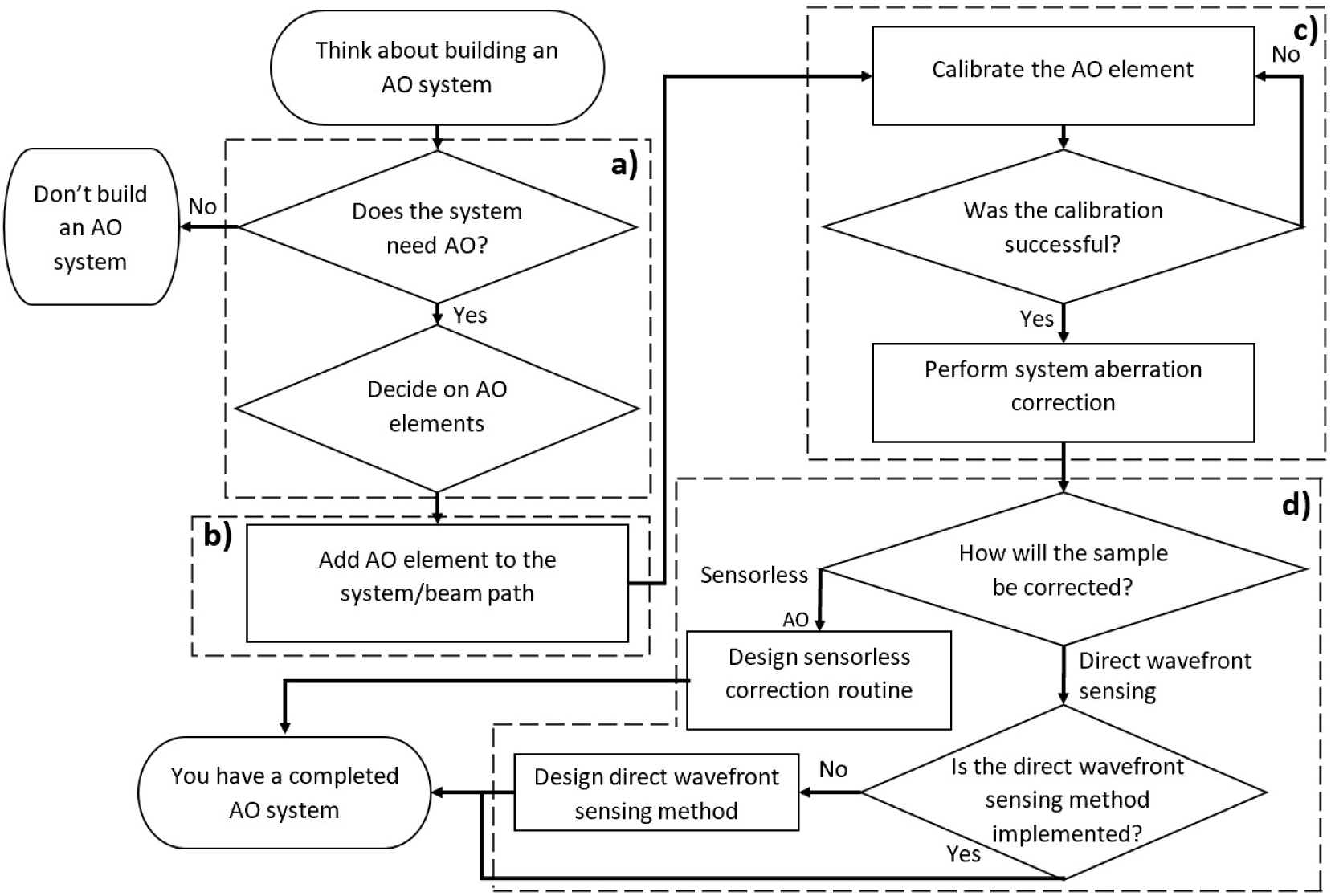
Flowchart depicting the general process for building a system utilising AO. a) *System Design Phase*. User decides whether the system needs AO and, if so, what type. b) *Installation Phase*. The chosen AO elements are added to the system. c) *Set-up Phase*. The chosen AO element is calibrated. Typically this involves mapping the variable elements of the AO element (e.g. deformable mirror actuators) to a useful set of basis functions which represent optical aberrations. d) *Sample Correction Phase* Here the user designs the methods to be used for correcting their desired sample.

1. *System Design Phase*: A potential user should consider the needs of their imaging modality, system constraints, desired sample types before deciding on the appropriate AO element to implement.
2. *Installation Phase*: The user installs the chosen AO element into their beam path.
3. *Set-up Phase*: The AO element is calibrated to correct for optical aberrations. This calibration is checked and the system aberrations are corrected.
4. *Sample Correction Phase*: The sample correction routine is designed. This will typically fall into one of two categories; sensorless AO or direct wavefront sensing.

Microscope-AOtools does not contain methods relevant to the *System Design* or *Installation* phases, although resources do exist to aid with these[14, 15]. Utilising Microscope-AOtools requires that the adaptive element that the user has decided on is a Python-Microscope compatible device and that the user has some kind of wavefront sensor installed (e.g. interferometer, Shack-Hartmann sensor). However, There are no other requirements for using Microscope-AOtools. Microscope-AOtools provides all the methods necessary for the *Set-up* and *Sample Correction* phases, easing the development of AO enabled microscopes.

## 0.3 Methods

### Calibration

The general principle of aberration correction is to measure the overall aberration of the optical wavefront and apply the opposite phase deformation to the adaptive element. An AO device is generally composed of *N* variable components i.e. a deformable mirror has *N* actuators which control the shape of the mirror surface. Measuring the phase wavefront directly, these *N* variable components can shape the adaptive element to correct for the local phase distortions. However, in many AO devices these components are coupled, have non-linear responses, or other non-ideal behaviours, as is the case for one of the most common AO devices, the continuous membrane deformable mirror[16]. If the phase wavefront is not directly observable, attempting aberration correction by varying individual components of the AO device is prohibitively difficult. Therefore, the use of adaptive elements requires a map between the variable components and the aberrations we wish to correct, allowing the whole of the adaptive element can be configured at once to correct for phase distortions. Constructing this map is the calibration process.

Consider a continuous membrane deformable mirror as our adaptive element. Assuming that the overall mirror shape is the linear superposition of all the individual actuator deflections, we can define the overall mirror shape, *S*(*x, y*) as:

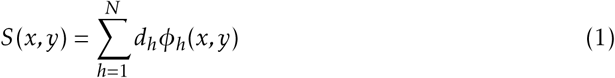

Where *S*(*x, y*) is the change in the deformable mirror shape from its original position, *d*_*h*_ is the *h*-th actuator control signal (an arbitrary value related to applied voltage which determines the position of the *h*-th actuator in its overall movement range) and *ϕ*_*h*_(*x, y*) is the *h*-th influence function. So called as they describe how the elements of the device influence the phase wavefront. We can convert this set of basis functions to a different basis set. An obvious alternative basis set is the Zernike polynomials since they are defined on the unit circle, orthogonal and the wavefront distortion can be well approximated by the linear addition of a limited number of Zernike polynomials[17, 18]. Describing *ϕ*_*h*_(*x, y*) in terms of Zernike polynomials we obtain:

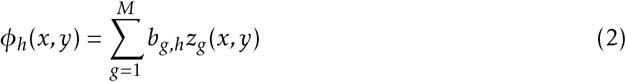

Where *b*_*g,h*_ is the coefficient corresponding to the *k*-th Zernike polynomial due to *d*_*h*_, the *h*-th actuator control signal. This leads to:

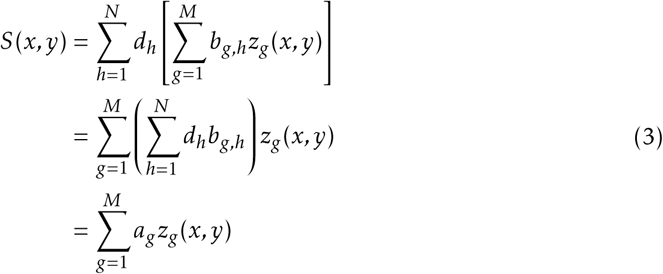

Where the new Zernike coefficients, *a*_*g*_, are defined as:

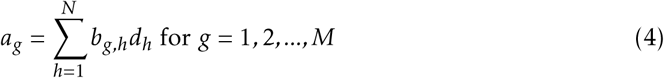

Converting this to a matrix form yields:

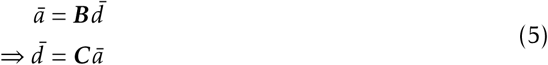

Where 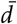 is a length *N* vector of the actuator control signals, *ā* is the length *M* vector of the Zernike polynomial amplitudes and *B* is the *M* × *N* matrix representing the response characteristics of the deformable mirror. However, we actually want its inverse, *B*^−1^ = *C*, otherwise called the control matrix, in order to convert from Zernike polynomial amplitudes to actuator control signals.

Microscope-AOtools implements an automated calibration routine to obtain *C*. Each actuator is moved through *p* set positions and a wavefront is extracted. The wavefront is then decomposed into *M* Zernike modes[19]. A row vector *z* is obtained for each actuator position containing the computed Zernike mode amplitudes:

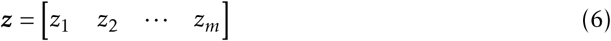

Where the *g*-th element is the amplitude of the *g*-th Zernike mode. By collecting the row vectors of each position for the *h*-th actuator we can obtain:

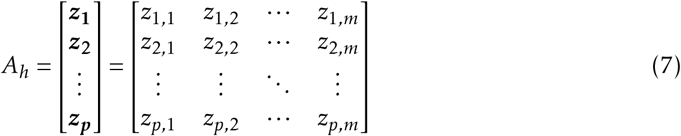

Linear regression to each column, [*z*_1,*i*_ *z*_2,*i*_ … *z _p,i_*]^*T*^, yields the response characteristics between the *h*-th actuator’s position and the *g*-th Zernike mode, *b*_*g,h*_. In this way, we construct ***B*** and then calculate ***C***. Figure 2 shows a flowchart of this process as implemented in Microscope-AOtools.

**Figure 2:**
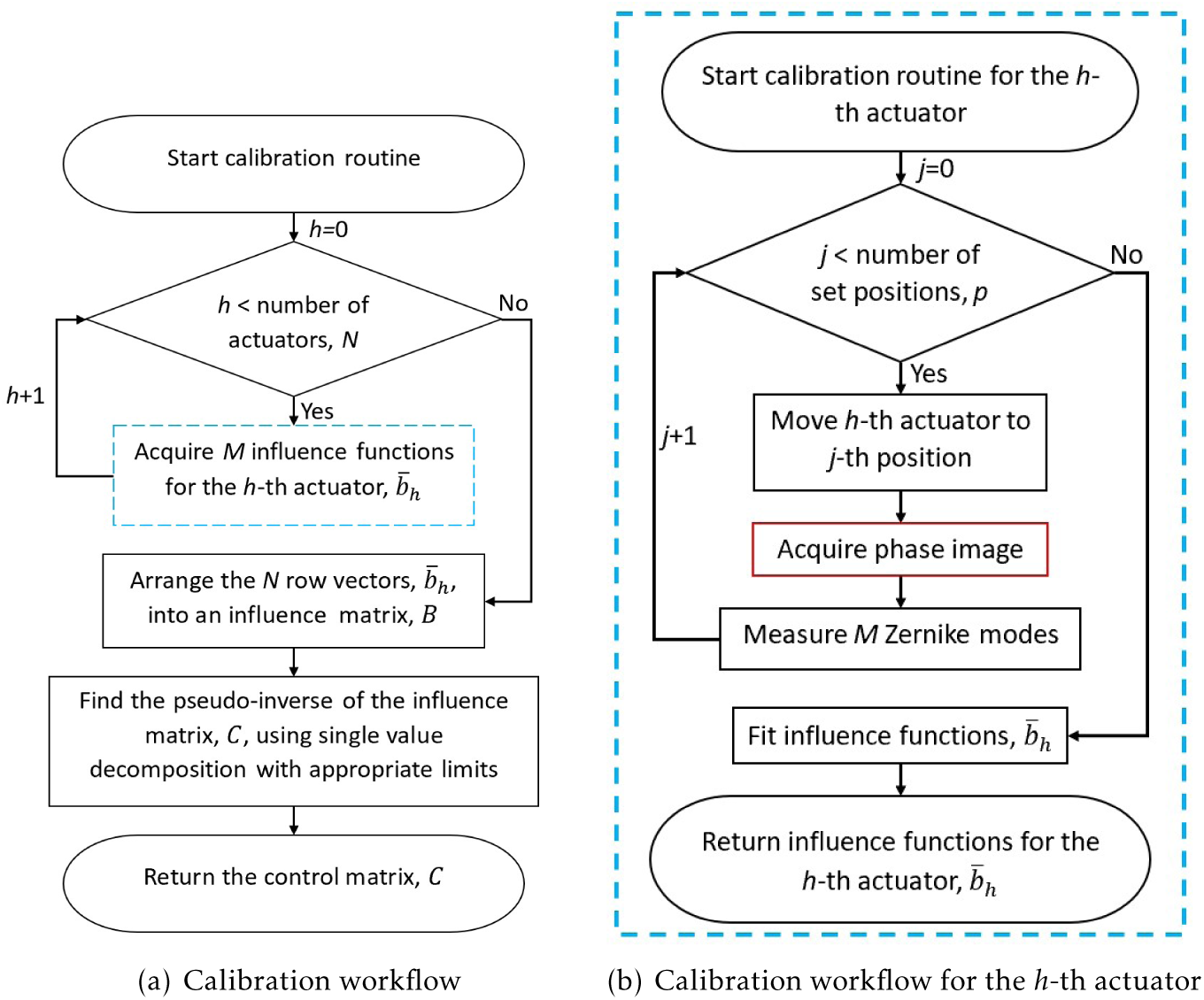
(a) Flowchart depicting the generalised calibration routine implemented in Microscope-AOtools (b)Flowchart depicting the process for calibrating the *h*-th actuator of the deformable mirror, the dashed blue process in (a). The influence functions returned are *b*_*g,h*_ described in Equation 2. This process is performed for each of *N* actuators and used to obtain *C* described in Equation 5.

In general, ***B*** is singular, or near singular, and therefore has no true inverse. So we must use a pseudo-inverse, calculated using single value decomposition (SVD). Actuators that have little influence on particular Zernike modes will have small values in the *B* matrix. These small influence values occur due to a combination of the actuators being in physical positions where they have limited influence over certain Zernike modes and noise, which leads to small perturbations in the measured Zernike mode amplitudes which are unrelated to the actuator movement. A control matrix calculated without thresholding out these small values before inversion will quickly lead to a saturation of the deformable mirror actuators (i.e. actuators at their maximum stroke length) when corrections are calculated[20]. This occurs because small values in ***B*** become large values in the ***C***, which results in large actuator signals, *d*_*i*_, even at low Zernike mode amplitudes for certain actuators which try to correct modes which they have minimal influence over. Therefore, the calibration method incorporates a threshold by default and the exact threshold can be varied by experienced users.

Typically, a calibration routine is designed around a particular wavefront sensing method for one specific adaptive element and requires redesign for any new wavefront sensing technique. Microscope-AOtools does not make an assumption about the wavefront sensing technique used. Instead a raw image from the wavefront sensor is passed to one of a suite of phase acquisition methods and a phase image is returned. Which method in the suite is used is defined at start-up and can be changed by the user at any time.

Although the calibration routine has been defined in terms of a deformable mirror and its actuators, in principle it can be used to create a control matrix for an arbitrary adaptive element with *N* variable components (i.e. degrees of freedom). Microscope-AOtools queries the Python-Microscope device to discover the number of variable elements and is therefore able to calibrate for an arbitrary AO device with *N* degrees of freedom. This attribute is fetched from the device and used by Microscope-AOtools to calibrate every actuator/variable element for any arbitrary adaptive element with *N* degrees of freedom. By constructing the calibration workflow in this generalised manner Microscope-AOtools can be used on any arbitrary Python Microscope compatible adaptive element with any wavefront sensing technique.

### Characterisation

Feedback on the quality of the calibration process is essential. Ideally, once the adaptive element is calibrated we have a linear map which allows known quantities of Zernike modes to be applied exactly. This linear map is never exact in practice due to a range of issues such as, the fact that some parameters, like the number of steps used to calibrate each actuator and the threshold used in the SVD pseudo-inversion, are chosen empirically. Additionally, the approximate nature of the pseudo-inverse and discretisation errors (due to discrete sampling of a continuous functions) in the measuring of Zernike modes influence the quality of the linear map. It is therefore necessary to have some measure of how well the adaptive element is able to recreate desired Zernike modes. This process is called characterisation. It involves applying a fixed amplitude of single Zernike mode to the adaptive element, measuring the Zernike modes present in the wavefront and comparing to that applied. An automated implementation of this process is present in Microscope-AOtools with the results returned to the user for interrogation. Figure 3 shows a flowchart of this method in Microscope-AOtools.

**Figure 3:**
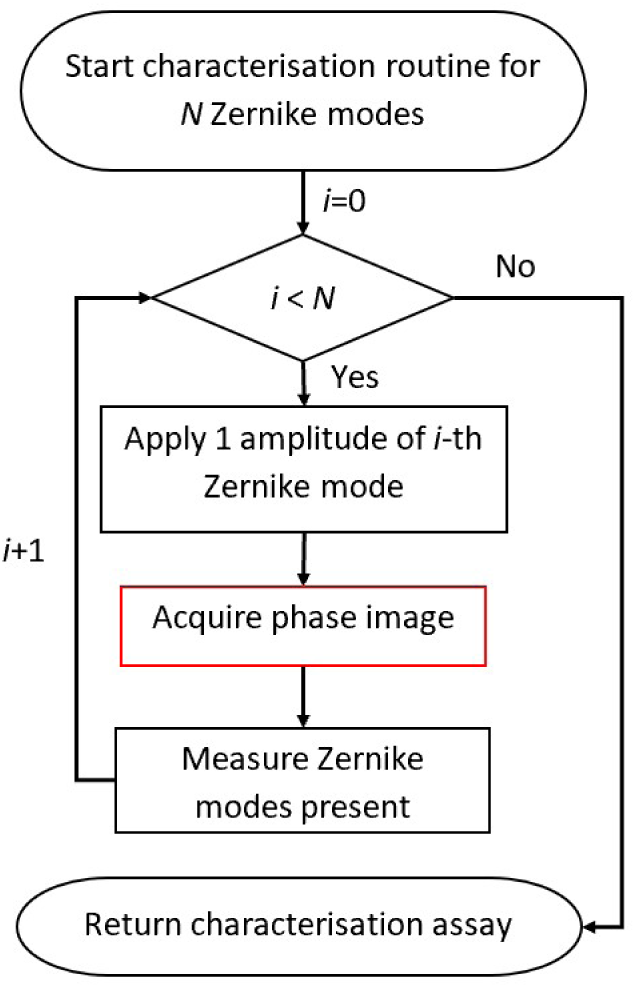
Flowchart depicting the process for characterising an adaptive element as implemented in Microscope-AOtools

In an ideal situation, where the control matrix provided a perfect linear map from Zernike mode amplitudes to the adaptive elements degrees of freedom, a characterisation assay like Figure 4(a) is expected, where only the Zernike mode applied has a none zero measured amplitude. In practice, the adaptive element is better at recreating particular Zernike modes and some Zernike mode coupling is observed, i.e. modes which were not applied to the adaptive element have measurable amplitude. This leads to characterisation plots like Figure 4(b). Here we present a characterisation assay obtained for an Alpao-69 actuator deformable mirror.

**Figure 4:**
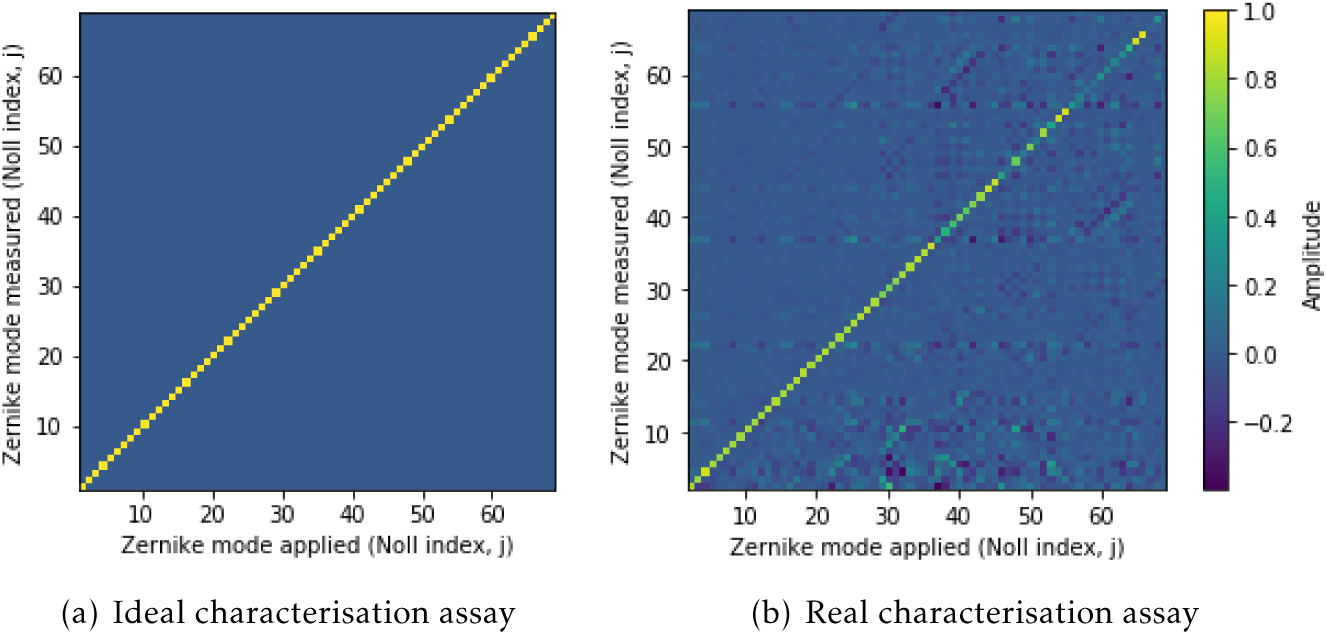
(a) An ideal characterisation assay, measuring the recreation accuracy of 68 Zernike modes with applied amplitude of 1 for each (b) A realistic characterisation assay obtained from a calibrated Alpao-69 actuator deformable mirror, measuring the recreation accuracy of 68 Zernike modes with applied amplitude of 1 for each

From this characterisation assay, various measures of calibration accuracy can be extracted, principally the amplitude of the applied Zernike mode and the amplitudes of the other, coupled, Zernike modes. Clearly not all Zernike modes are recreated equally well and different modes exhibit varying degrees of mode coupling. As previously mentioned, this arises from mathematical approximations, computational errors and the physical characteristics of the adaptive element. Microscope-AOtools provides the tools to assess the accuracy of Zernike mode recreation, which can be used to inform which modes should be included in the aberration correction algorithms.

The characterisation routine relies on the same generalised phase acquisition method used in the calibration workflow. Recall that this is a user selected method from a suite of phase acquisition methods. The number of Zernike modes assessed, *N*, is the number of modes that have been measured in the calibration step by default, but this can be varied by the user. Once again this preserves generalisability and allows the characterisation method to be used on any arbitrary adaptive element, calibrated for any number of modes and utilising any desired wavefront sensing technique.

### System Aberration correction

Microscope-AOtools implements a method for correcting the system aberrations via direct wavefront sensing, designed to be used after calibration and characterisation. The workflow is shown in Figure 5. The wavefront is obtained through whatever direct wavefront sensing method has been implemented and selected, a number of Zernike modes determined by the user are fitted to the wavefront, an equal and opposite magnitude of these modes are applied to the adaptive element. The RMS wavefront error is then obtained. This process repeats until *N* iterations have been performed or the RMS wavefront error is below a user defined error threshold, *δ*.

**Figure 5:**
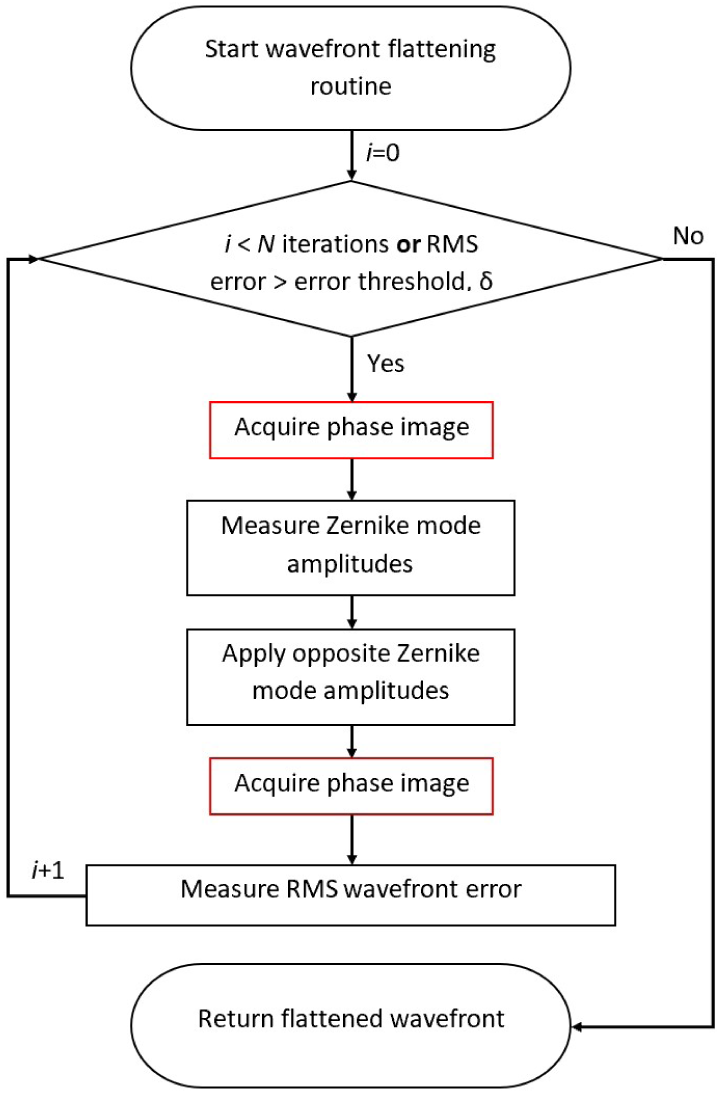
Flowchart depicting the process for flattening directly measured wavefront as implemented in Microscope-AOtools

It is necessary to perform this process for a number of iterations to ensure the optimal wavefront is obtained due to the limitations of Zernike mode recreation accuracy discussed previously. Figure 6 shows the results of one such wavefront correction, performed using the same Alpao-69 actuator deformable mirror as before. The wavefront was obtained by interferometry and Zernike modes 5-29 (using Noll indices). These modes were selected using the characterisation assay presented in Figure 4 and were corrected over 20 iterations. Figure 6(c) shows the Zernike mode amplitudes before and after correction. Both the Zernike mode amplitudes and RMS wavefront error show a significant improvement in wavefront quality.

**Figure 6:**
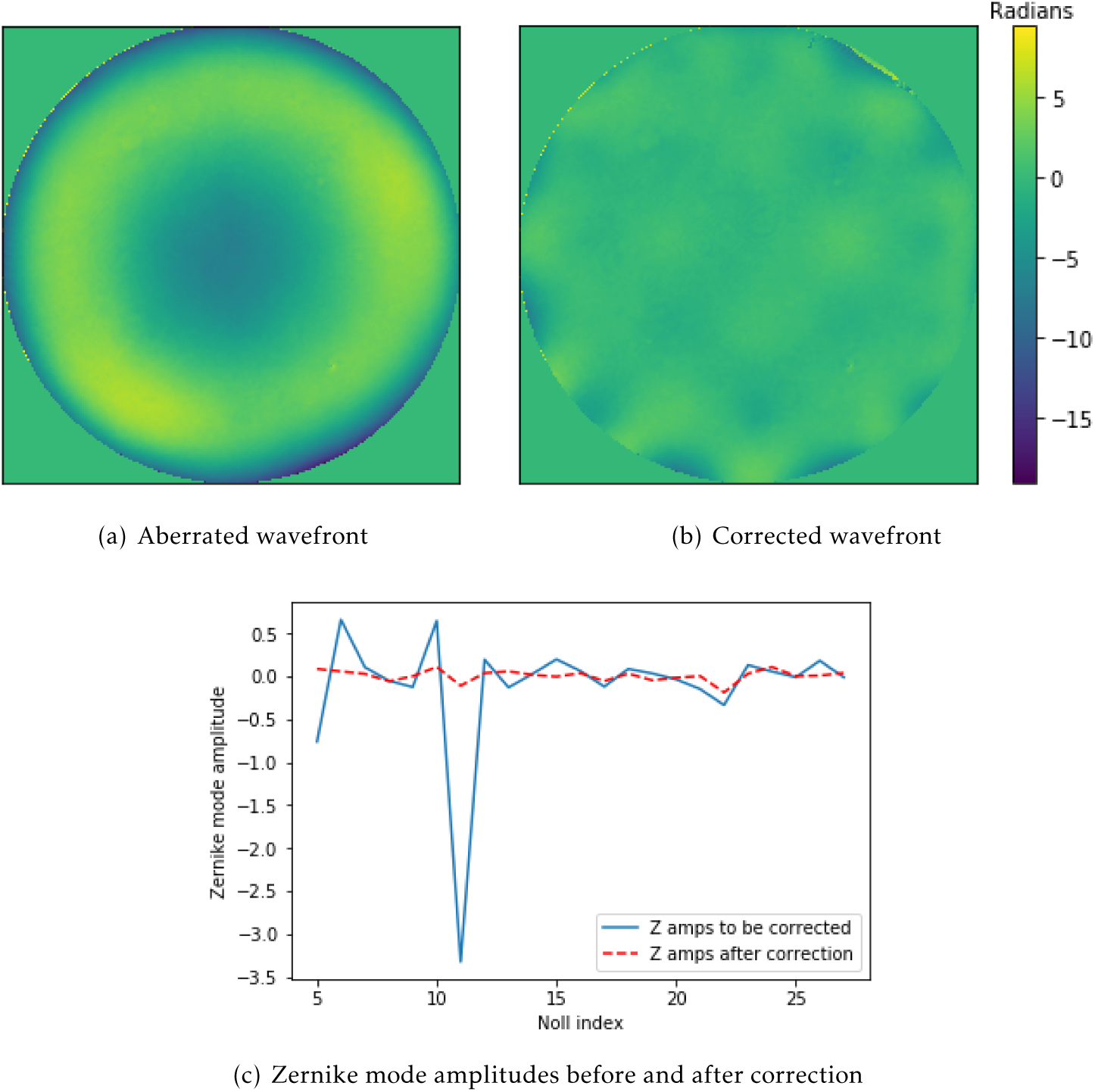
(a) An aberrated wavefront. RMS wavefront error = 3.818 radians (b) A wave-front after 20 iterations of correction. RMS wavefront error = 0.986 radians. RMS wave-front error of the central 95% of the phase wavefront = 0.712 radians(c) The Zernike modes measured in the aberrated (red) and corrected (blue) wavefronts (a) – (b) are all presented on the same colour scale (in radians of 543 nm HeNe laser) and were obtained via interferometry

For system aberration corrections it may be preferable to set a minimum wavefront error and continue to iterate until this threshold is reached. However a user may wish to only spend *N* iterations correcting the wavefront. Microscope-AOtools implements both options to ensure generalisability and which criteria is used can be set by the user. As with the calibration and characterisation methods, the wavefront flattening routine relies on a user selected phase acquisition technique from the suite of implemented methods. This ensures that the applicability of Microscope-AOtools to any adaptive element with any wavefront sensing technique is preserved throughout all the set-up methods.

## 0.4 Sample Correction Methods

### Direct Wavefront Sensing Correction

Performing AO correction for biological samples by directly measuring the phase wave-front has been well documented. In many cases the phase acquisition methods used are the same as those a user might implement to calibrate the adaptive element, usually a Shack-Hartmann wavefront sensor[21, 22, 23]. Occasionally, other methods are used[24]. Fortunately, since the wavefront correction workflow shown in Figure 5 has been kept generalised to allow any phase wavefront sensing technique to be used, the same workflow can be used for correcting the sample induced aberration as the system aberrations. Critically, the phase acquisition method, number of iterations and error threshold do not have to be the same in both processes. This is important since correcting for sample induced aberrations adds additional limitations. Biological samples can suffer damage when exposed to excessive light (phototoxicity). Repeated activation can also cause chemical alteration to fluorophores leading to inactivation (photobleaching). Microscope-AOtools is designed so a user can correct for as many iterations as required until a desired wavefront flatness is achieved or for exactly *N* iterations. The former is designed for system aberration correction, while the latter is designed for correcting sample induced aberrations in order to limit phototoxicity and photobleaching.

### Sensorless Correction

In many biological applications, direct wavefront sensing is not possible and so we rely on sensorless techniques to determine the best correction to apply. The generalised methodology for this is shown in Figure 7 for a biological specimen. Some metric, *S*, which gives a useful measure of the image quality is chosen. This metric should be a numerical value and should increase to a global maximum as the aberrations present decrease. Often these metrics are related to common measures of image quality, such as sharpness or contrast. For each Zernike mode, *Z*_*i*_, a number of amplitudes of that mode, *a*_*j*_, are applied and an image of the sample obtained. The quality of each image, *S*_*j*_ is calculated. Assuming that *S* is a function of the Zernike mode amplitude applied, fitting a Gaussian function to the *S*_*j*_ values yields a Zernike mode amplitude, *a*_*max*_, which theoretically yields the best image quality, *S*_*max*_. The complexity of sensorless AO correction lies in selecting the most appropriate image quality metric. There have been numerous metrics developed which have been shown to be effective on certain sample types or imaging modalities[7, 25, 26, 27].

**Figure 7:**
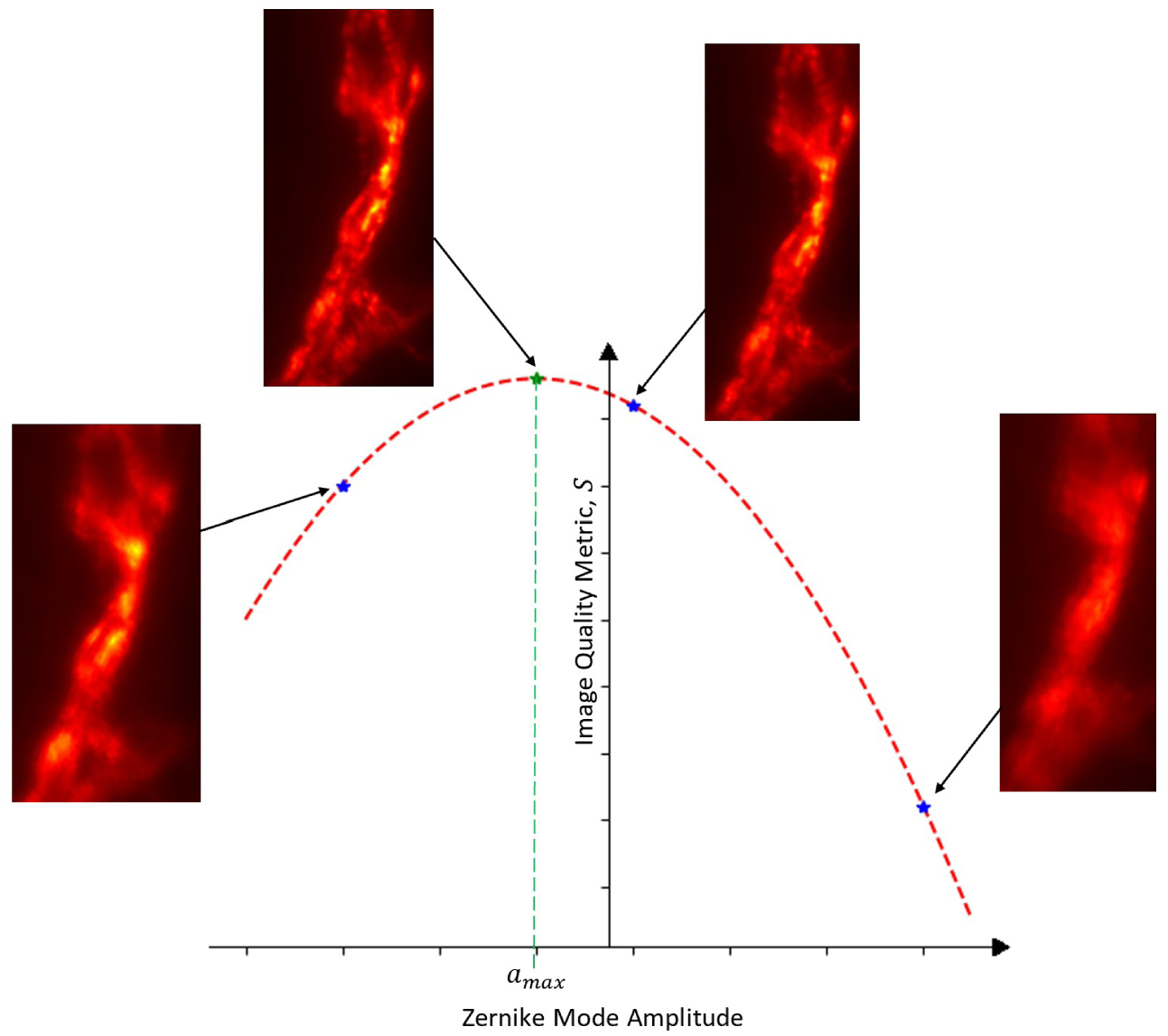
Principle of sensorless AO correction. The inset images are Drosophila Neuro-muscular Junction (NMJ). For each Zernike mode, *Z*_*i*_, NMJ images were acquired for different amplitudes of the *i*-th Zernike mode. A value of the image quality metric, *S*, is obtained for each (blue dots). A Gaussian function is then fitted to these values and the amplitude, *a* corresponding to the maximum image quality, *S*_*max*_, is obtained (green dot). The inset figure for the green spot shows the NMJ image acquired after the correction for the *i*-th Zernike mode was applied

An automated sensorless AO routine is not implemented in Microscope-AOtools as this is outside its scope as it would require giving Microscope-AOtools control over the complete imaging system. Python-Microscope, which Microscope-AOtools uses, already fulfils this role. There are three options for sensorless correction workflows. The first, Figure 8(a), an amplitude, *a*_*j*_, of the *i*-th Zernike mode is applied, an image of the sample is taken and the image quality metric is evaluated. This process is repeated for *M* measurements and then the Zernike mode amplitude corresponding to the maximum image quality, *a*_*max*_, is calculated and applied. This process is repeated for *N* Zernike modes. The second, Figure 8(b), is broadly similar with the exception that the image quality metric for the *M* images of the current Zernike mode are measured after they have all been acquired rather than as soon as each image is acquired. The final workflow option, Figure 8(c), differs further by not applying each Zernike mode correction sequentially but rather calculating the image quality metrics for all *NM* images at the end of the imaging routine and calculating the correction to be applied for all *N* Zernike modes simultaneously.

**Figure 8:**
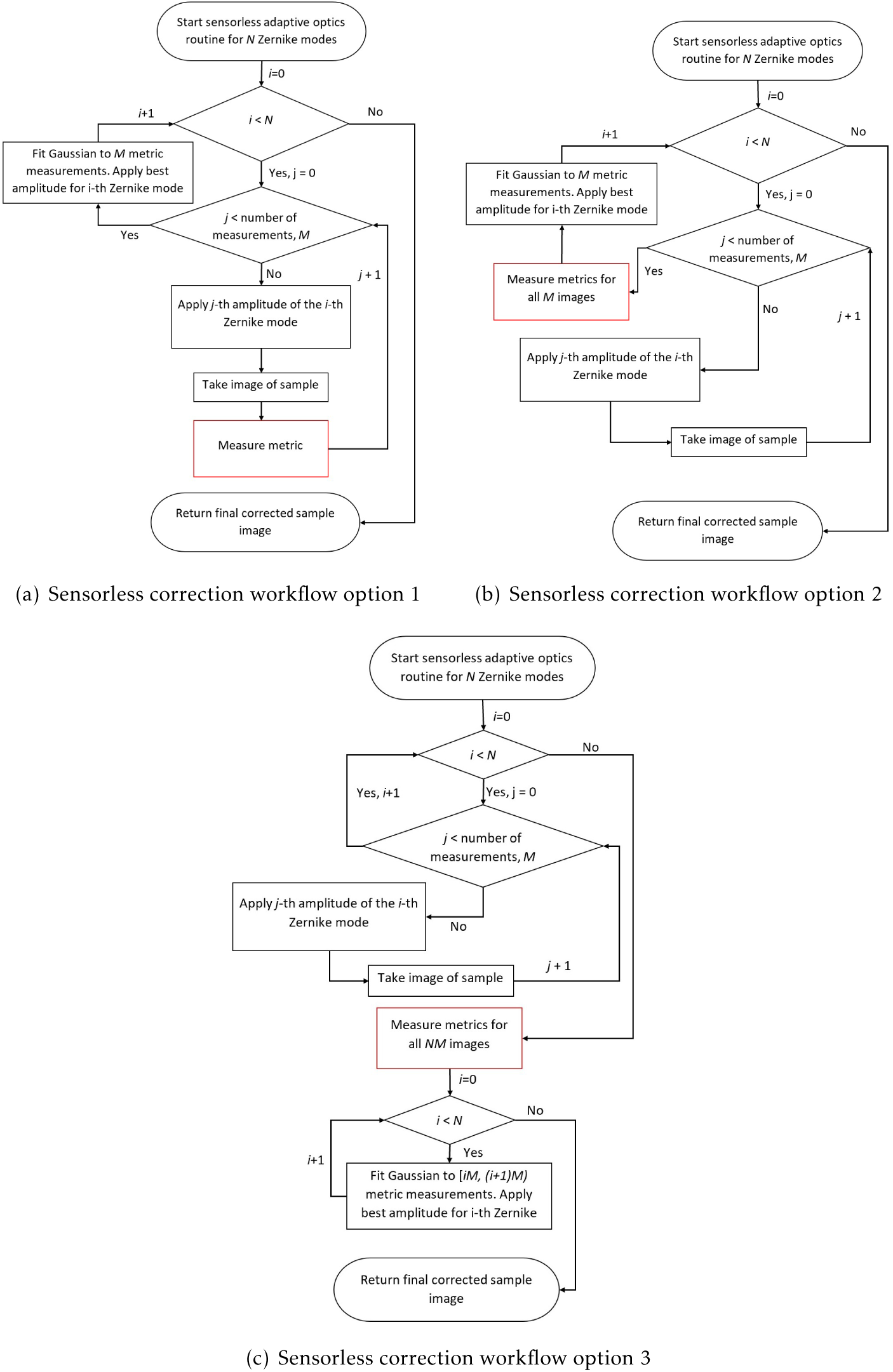
Flowcharts depicting the sensorless correction routine options (a) An image for each amplitude of the *i*-th Zernike mode is taken and the image quality metric is immediately evaluated. Once all the images for the *i*-th Zernike mode have been taken, the best Zernike amplitude is found as described in Figure 7 and applied (b) All *M* images are taken, then the quality metric is obtained for all *M* images, the best Zernike amplitude is found and applied (c) All the images for all the *N* Zernike modes are obtained with no correction applied in between modes. The image quality metric then measured for every image and the best amplitude for each Zernike mode is found. The correction for all modes is applied simultaneously at the end of the workflow.

Similar to the set-up methods, a sensorless AO workflow is typically developed for a specific sample type or imaging modality and any change to these specifics requires redesigning the entire workflow. The only significant difference between these implementations will be the image quality metric used. Microscope-AOtools makes no assumption about the desired image quality metric. Instead a raw image is passed to one of a suite of image quality metrics and a metric value is returned. The image quality metric used can be easily changed allowing the user to select a quality metric optimised for their sample type and imaging modality. Microscope-AOtools also implements the methods necessary for all three of the workflows shown in Figure 8 allowing a user to select their preferred workflow.

### IsoSense

Anisotropies in the sample structure can bias the corrections towards improving the image quality in a non-uniform manner. There has recently been a technique developed to overcome this issue: IsoSense, which relies on producing spatially structured light in order to fill empty sections of the image Fourier spectrum.[28] IsoSense is designed to be used in structured illumination microscopy (SIM) setups since they often incorporate spatial light modulators (SLM) as high-speed, dynamic diffraction gratings and SIM is particularly sensitive to Fourier space anisotropies.

Microscope-AOtools incorporates the methods necessary to implement IsoSense. Figure 9 shows both the structured illumination pattern used for IsoSense, which is applied to an SLM, and the location of the beams in Fourier space. The illumination pattern shown in Figure 9(a) is the inverse Fourier transform of the 4-beam interference pattern in Figure 9(b). The location of these beams are: 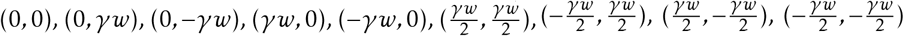. *w* is the Abbe diffraction limit and *γ* is a user defined fill fraction. This fill fraction controls the positions of the beams in the interference pattern and hence the region of the Fourier spectrum which will be enhanced over normal illumination. Placing these beams is a skill for more advanced users, however the implementation in Microscope-AOtools has a sensible default and advanced users can improve their AO correction further by manipulating this if needed.

**Figure 9:**
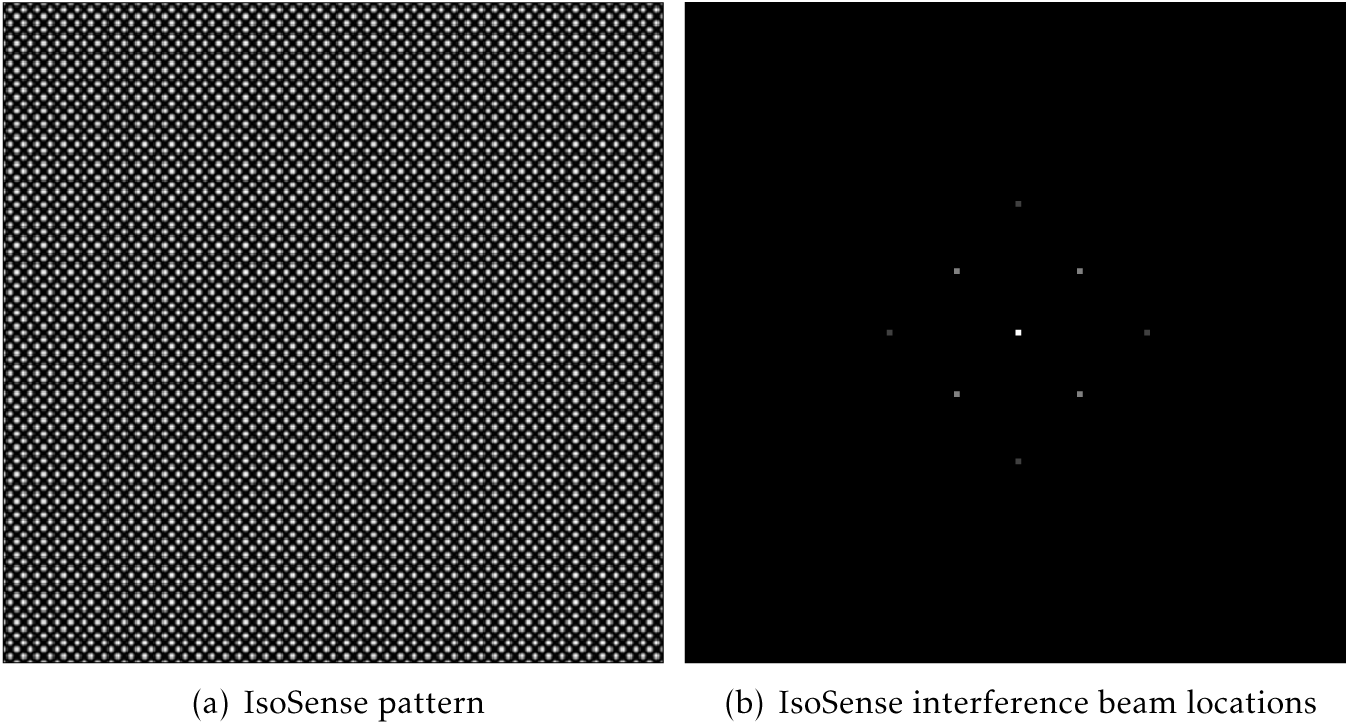
(a) A simulated IsoSense pattern created with a 4-beam interference. A pattern similar to this is applied to the SLM (b) A diagram of a 4 beam interference pattern in Fourier space.

## 0.5 Future Expansion

So far we have discussed the specific methods implemented in Microscope-AOtools. The Set-up and Sample Correction methods rely on suites of techniques, wavefront sensing and image quality metric assessment respectively. These are designed to be easily extensible by users as new techniques are developed. The functions defining the existing wavefront sensing and image quality metric assessment techniques are stored in separate files. Which wavefront sensing technique will be used is an attribute of the highest level of the code hierarchy and is used to select a wavefront sensing technique from the unwrapping method dictionary. Similarly, which image quality metric assessment will be used is an attribute of a lower level of the code hierarchy and is used to select a wavefront sensing technique from the dictionary of image quality metrics. A detailed guide of where the appropriate classes and dictionaries are located and how to add new wavefront sensing and image quality metric assessment techniques is included in the *README.md* file for Microscope-AOtools. Briefly, these suites are composed of functions with a defined set of input and output variables. A user creates a new wavefront sensing or image quality metric assessment functions with the input and outputs defined in the *README.md*, adds this function to the correct file and then adds the function option to the correct suite dictionary.

## 0.6 Discussion

Microscope-AOtools has been designed so a user can take an adaptive element in an abitrary set-up, calibrate the adaptive element and use it on any sample type in an range of imaging modalities. Since Microscope-AOtools leverages Python-Microscope it already supports a number of adaptive elements, mostly deformable mirrors, which will expand as hardware support in Python-Microscope expands. Adding new devices to Python-Microscope is relatively simple. Refer to Python-Microscope (https://www.python-microscope.org/) for more details. Microscope-AOtools only requires that the adaptive element be a Python-Microscope device which has an attribute *n_actuators* which defines the number of variable components of the device.

The process of setting up an adaptive element requires a wavefront sensor to observe the shape of the phase wavefront and calibrate how the variable components of the adaptive element affect this wavefront. By designing the set-up methods in Microscope-AOtools to accept any method from a suite of wavefront sensing techniques, Microscope-AOtools is both generalised and easily extensible. If the desired wavefront sensing technique is not already incorporated then a user only has to add the function necessary to perform the wavefront sensing step rather than to reimplement the set-up methods in their entirety. Microscope-AOtools is further generalised as it allows for the control matrix to be acquired by some external method and then set in Microscope-AOtools with the *set_controlMatrix* method. This ensures that a user with an existing calibration routine wishing to access the sensorless AO methods in Microscope-AOtools can do so without having to repeat work they have already performed. It also allows control matrices acquired from routines using different phase acquisition techniques to be compared. Characterisation assays can be acquired for each method’s control matrix and the accuracy of the Zernike mode recreation compared.

The generalised nature of Microscope-AOtools continues into the Sample Correction methods. By allowing the user to swap between wavefront sensing technique, Microscope-AOtools already possesses all the methods necessary for performing sample correction using direct wavefront sensing, provided the wavefront sensing technique is already included in the suite of methods. Similarly Microscope-AOtools utilises a suite of image quality metrics suited to different sample types and imaging modalities. A user can select a pre-existing metric well suited to their application. If no appropriate metric currently exists an new one can easily be implemented and added to the suite of metrics. Once implemented it can be used in any of the sensorless AO analysis methods outlined in Figure 7. Furthermore, Microscope-AOtools allows for Zernike mode amplitudes to be set directly with the *set_phase* method. This means that if a user has an offline analysis technique, such as a machine learning approach, Microscope-AOtools can be used to calibrate the deformable mirror, the sample induced calculations are performed offline and then the appropriate correction applied through Microscope-AOtools.

Microscope-AOtools is free and open-source. It is intended to be a resource for the microscopy community at large and it is designed to minimise the time and effort spent replicating work other AO users have already done. As Microscope-AOtools acquires a larger base of users, some adding their own wavefront sensing techniques or image quality metrics to expand the existing suites, future and existing users will have a wider array of usability options, accelerating the adoption of novel techniques by the microscopy community and lowering the barrier to entry to set-up an AO system.

Beyond the open-ended task of expanding the existing suite of phase acquisition techniques and image quality metrics, there are a number of future developments that could be made to Microscope-AOtools. There does not currently exist a universal image quality metric, although strides have been made in that direction.[29] Image quality metrics attempt to assign a numerical value for how ‘good’ an image is, but what makes a ‘good’ image varies between imaging modalities, sample type and even users. Most metrics pick some aspect of the image deemed to be significant (e.g. contrast, sharpness, maximum intensity, etc) and maximise it. Since Microscope-AOtools has access to multiple image quality metrics, one development would be designing a sensorless AO routine which measures multiple image qualities simultaneously, assigns some weight to each metric measurement and maximises the image quality based on several criteria.

## 0.7 Conclusion

For some time, there has been a call for a robust, generalised implementation for AO. Such an implementation should incorporate all the methods needed to setup and operate an AO element for a range of imaging modalities and sample types. Microscope-AOtools includes methods for calibration, direct wavefront and sensorless correction. In particular, it already incorporates several image quality metrics suited to sensorless correction in a number of different imaging modalities. It also includes a characterisation method for assessing the accuracy of the calibration step. It has also been designed in a modular manner allowing for new wavefront sensing techniques and image quality metrics to be added with minimal disruption to the rest of the workflows and, therefore, minimal work duplication. With time and community support, such an implementation has scope to go beyond its current state of “generalised software implementation” and become a universal software implementation for AO.

## Software availability

1. microscope-aotools code repository: https://github.com/MicronOxford/microscope-aotools
2. Software license: GNU General Public License version 3 or any later version (https://www.gnu.org/licenses/gpl-3.0.en.html)

## Acknowledgements

The authors would like to thank Mick Philips, Mantas Žurauskas, Jacopo Antonello and David Pinto for their helpful comments and suggestions during the development of Microscope-AOtools

## References

[1] Michael Schwertner, Martin J Booth, and Tony Wilson. “Characterizing specimen induced aberrations for high NA adaptive optical microscopy”. In: Optics express 12.26 (2004), pp. 6540–6552.

[2] Michael Schwertner et al. “Measurement of specimen-induced aberrations of biological samples using phase stepping interferometry”. In: Journal of microscopy 213.1 (2004), pp. 11–19.

[3] Martin J Booth. “Adaptive optics in microscopy”. In: Philosophical Transactions of the Royal Society of London A: Mathematical, Physical and Engineering Sciences 365.1861 (2007), pp. 2829–2843.

[4] James C Wyant and Katherine Creath. “Basic wavefront aberration theory for optical metrology”. In: Applied optics and optical engineering 11.part 2 (1992), pp. 28–39.

[5] Martin J Booth. “Adaptive optical microscopy: the ongoing quest for a perfect image”. In: Light: Science & Applications 3.4 (2014), e165.

[6] John M Girkin, Simon Poland, and Amanda J Wright. “Adaptive optics for deeper imaging of biological samples”. In: Current opinion in biotechnology 20.1 (2009), pp. 106–110.

[7] Daniel Burke et al. “Adaptive optics correction of specimen-induced aberrations in single-molecule switching microscopy”. In: Optica 2.2 (2015), pp. 177–185.

[8] Daniel E Milkie, Eric Betzig, and Na Ji. “Pupil-segmentation-based adaptive optical microscopy with full-pupil illumination”. In: Optics letters 36.21 (2011), pp. 4206–4208.

[9] Kai Wang et al. “Rapid adaptive optical recovery of optimal resolution over large volumes”. In: nature methods 11.6 (2014), pp. 625–628.

[10] Peter Kner et al. “Adaptive optics in wide-field microscopy”. In: MEMS Adaptive Optics V. Vol. 7931. International Society for Optics and Photonics. 2011, 79310K.

[11] Dean Wilding et al. “Adaptive illumination based on direct wavefront sensing in a light-sheet fluorescence microscope”. In: Optics express 24.22 (2016), pp. 24896–24906.

[12] Cristina Rodríguez and Na Ji. “Adaptive optical microscopy for neurobiology”. In: Current opinion in neurobiology 50 (2018). AO application in two photon neurobiology imaging. Contains lines in the discussion & conclusion which call for “methods that can provide aberration correction for a variety of specimens, in a fast and accurate fashion, which, once implemented, is robust and simple to use.”, pp. 83–91.

[13] Na Ji. “Adaptive optical fluorescence microscopy”. In: Nature methods 14.4 (2017), p. 374.

[14] M. J. Booth. A basic introduction to adaptive optics for microscopy. Oct. 2019. doi: 10.5281/zenodo.3471043. URL: https://doi.org/10.5281/zenodo.3471043.

[15] I Dobbie, N Hall, and DMS Pinto. “BeamDelta: simple alignment tool for optical systems”. In: Wellcome Open Research 4 (2019).

[16] Lijun Zhu et al. “Adaptive control of a micromachined continuous-membrane deformable mirror for aberration compensation”. In: Appl. Opt. 38.1 (1999), pp. 168–176. doi: 10.1364/AO.38.000168. URL: http://ao.osa.org/abstract.cfm?URI=ao-38-1-168.

[17] Zernike by F. “Diffraction theory of the cutting process and its improved form, the phase contrast method”. In: Physica 1.7-12 (1934), pp. 689–704.

[18] Robert J Noll. “Zernike polynomials and atmospheric turbulence”. In: JOsA 66.3 (1976), pp. 207–211.

[19] MJ Townson et al. “AOtools: a Python package for adaptive optics modelling and analysis”. In: Optics express 27.22 (2019), pp. 31316–31329.

[20] Martin Booth et al. “Methods for the characterization of deformable membrane mirrors”. In: Applied optics 44.24 (2005), pp. 5131–5139.

[21] Oscar Azucena et al. “Adaptive optics wide-field microscopy using direct wavefront sensing”. In: Optics letters 36.6 (2011), pp. 825–827.

[22] Xiaodong Tao et al. “Adaptive optics confocal microscopy using direct wavefront sensing”. In: Optics letters 36.7 (2011), pp. 1062–1064.

[23] Xiaodong Tao et al. “Live imaging using adaptive optics with fluorescent protein guide-stars”. In: Optics express 20.14 (2012), pp. 15969–15982.

[24] Saad A Rahman and Martin J Booth. “Direct wavefront sensing in adaptive optical microscopy using backscattered light”. In: Applied optics 52.22 (2013), pp. 5523–5532.

[25] Martin J Booth et al. “Adaptive aberration correction in a confocal microscope”. In: Proceedings of the National Academy of Sciences 99.9 (2002), pp. 5788–5792.

[26] JR Fienup and JJ Miller. “Aberration correction by maximizing generalized sharpness metrics”. In: JOSA A 20.4 (2003), pp. 609–620.

[27] Delphine Débarre et al. “Adaptive optics for structured illumination microscopy”. In: Optics express 16.13 (2008), pp. 9290–9305.

[28] Mantas Žurauskas et al. “IsoSense: frequency enhanced sensorless adaptive optics through structured illumination”. In: Optica 6.3 (2019), pp. 370–379.

[29] Jacopo Antonello et al. “Multi-scale sensorless adaptive optics: application to stimulated emission depletion microscopy”. In: Optics Express 28.11 (2020), pp. 16749–16763.

